# DeepCEF: A Deep Causal Estimation Framework for Complex Biological Systems Integrating Local Scores, Independence Tests, and Relation Attributes

**DOI:** 10.1101/2025.10.09.681515

**Authors:** Zhenjiang Fan, Mengrui Zhang, Summer Han

## Abstract

Causal relationship identification is a fundamental and complex research challenge that spans multiple disciplines, including biology, epidemiology, economics, and philosophy. Various scoring techniques and independence tests, such as local scores (e.g., Degenerate Gaussian (DG) and Bayesian Information Criterion (BIC)) and independence tests (e.g., Fisher’s Z), have been employed in causality estimation. However, these local scores often excel in specific data types or application areas but falter in others, limiting their ability to capture the complexity and heterogeneity of underlying causal mechanisms. For instance, a method may perform well on linear relationships or continuous variables but struggle with discrete variables or non-linear relationships.

Real-world observational datasets, particularly those generated in complex biological systems, often contain diverse data types and relationships, making it essential to develop a more comprehensive approach. To address this challenge, we propose a novel causal estimation framework that leverages the powerful classification capabilities of deep neural networks (DNNs) to identify causal patterns in pairwise relationships. Our framework integrates multiple local causality estimation scores, independence tests, and variable attributes, allowing it to capture a wide range of causal mechanisms.

To ensure the framework’s robustness and generalizability, we incorporate a diverse range of simulation data and 10 curated real-world datasets into the training procedure. Furthermore, our framework is designed to be extensible, enabling users to easily integrate their own data and additional scores and tests. Our validation results demonstrate that our framework outperforms existing methods in terms of estimation accuracy and precision on both simulation data and real-world biological datasets. By providing a more comprehensive and adaptable approach to causal relationship identification, our framework has the potential to advance research in various fields and improve our understanding of complex biological systems.

## Introduction

Causality research is generally divided into two main fields: causal discovery and causal inference (Nogueira et al. 2022, e1449). Causal inference focuses on estimating the effect of a treatment (or exposure) on an outcome. Unlike mere statistical correlations, causal inference seeks to establish whether changes in one variable *directly influence* another. Causal discovery, on the other hand, aims to uncover the underlying causal structure of a system from observational data. This involves identifying directed relationships (e.g., X→Y*X*→*Y*) rather than just dependencies. Common approaches include: 1) **Constraint-based methods** (e.g., the Peter and Clark (PC) algorithm (Spirtes, Peter, Glymour, and Scheines 2000) and Fast Causal Inference (FCI) algorithm (Spirtes, Peter L., Meek, and Richardson 2013) for latent confounders), which use conditional independence tests to prune possible causal graphs; 2) **Score-based methods** (e.g., Greedy Equivalence Search (GES) (Chickering 2002, 507–554)), which optimize a likelihood-based score over candidate graphs; 3) **Structural and functional causal models** (SCMs/FCMs), such as LiNGAM (Shimizu et al. 2006), which leverage functional assumptions (e.g., linearity or non-Gaussian noise) to resolve causal direction. Some constraint-based method, like PC and FCI, assume faithfulness while pruning edges from a fully connected graph (Ramsey, Zhang, and Spirtes 2006; Pearl 2009; Spirtes, Peter, Glymour, and Scheines 2000). This assumption states that the conditional independence relationships in the data are exactly those implied by the causal graph.

Causal inference and discovery are fundamental challenges in computational biology, where understanding the causal mechanisms underlying biological processes is essential for advancing research in areas such as gene regulation, disease pathways, and drug discovery. Unlike correlation-based approaches, which merely identify associations between variables, causal inference aims to uncover the direction and nature of causal relationships, enabling researchers to predict the effects of interventions and elucidate the mechanisms driving biological systems (Pearl 2009; Spirtes, Peter, Glymour, and Scheines 2000). This capability is particularly critical in complex biological systems, where interactions between genes, proteins, metabolites, and environmental factors give rise to intricate and often nonlinear relationships (Kitano 2002, 1662– 1664; Fan et al. 2023, gad044).

Despite significant advances in high-throughput technologies, such as genomics, transcriptomics, and proteomics, the identification of causal relationships from observational data remains a formidable task (Schadt et al. 2005, 710–717). Traditional methods for causal inference and discovery, such as local scoring techniques (e.g., BIC (Schwarz 1978, 461–464), DG (Andrews, Ramsey, and Cooper 2019, 4–21)) and independence tests (e.g., Fisher’s Z (Fisher, Ronald Aylmer 1921, 3–32), Conditional Mutual Information (Steeg and Galstyan a; Steeg and Galstyan b)), have been widely used to estimate causal relationships (Koller and Friedman 2009). However, these methods often excel only in specific data types or application domains. For instance, a method may perform well on linear relationships or continuous variables but struggle with discrete variables or nonlinear relationships (Peters, Janzing, and Sch olkopf 2017; Fan et al. 2023, gad044). This limitation hinders their ability to capture the complexity and heterogeneity of underlying causal mechanisms, particularly in real-world biological datasets, which often exhibit diverse data types, nonlinear interactions, and high dimensionality (Efron 2010). These complexities necessitate the development of more robust and comprehensive approaches to causal inference and discovery (Marbach et al. 2012, 796–804).

To address these challenges, we propose a novel deep-learning-based causal estimation framework (DeepCEF) that leverages the powerful classification capabilities of DNNs to identify causal patterns in pairwise relationships. Our framework integrates multiple local causality estimation scores, independence tests, and variable attributes, enabling it to capture a wide range of causal mechanisms. By incorporating a diverse range of simulation data and real-world biological datasets into the training procedure, we ensure the framework’s robustness and generalizability. Furthermore, the framework is designed to be extensible, allowing users to easily integrate their own data and additional scores or tests.

Our research advances the field of causal inference and discovery through four major contributions. First, we provide a comprehensive framework for causal inference and discovery that overcomes the limitations of traditional methods by leveraging the flexibility and power of DNNs. Second, we demonstrate the framework’s superior performance on both simulation data and real-world biological datasets, highlighting its ability to capture complex causal mechanisms. Third, we offer a user-friendly and extensible tool that can be readily applied to a wide range of biological problems, from gene regulatory network inference to drug discovery. Additionally, unlike constraint-based approaches, DeepCEF does not require the faithfulness assumption to hold for the inferred causal structure.

In the following sections, we describe the design and implementation of our framework, present validation results, and discuss its applications in computational biology. By providing a more comprehensive and adaptable approach to causal relationship identification, our framework has the potential to advance research in various fields and improve our understanding of complex biological systems.

## Methods

### 2.1 Theoretical Framework

This study aims to estimate the underlying causal relationships among variables from observational data, such as a data matrix. Given a dataset X ∈ R^n×d^, where n represents the number of observations and *d* represents the number of variables, the objective is to uncover the causal structure that governs the data-generating process (Figure 1.). This causal structure is represented as a directed acyclic graph (DAG) 𝒢= (V, E), where V is the set of nodes (variables) and E is the set of directed edges (causal relationships). **Figure 1**. provides an overview of the proposed framework. **Figure 1.D**. represents the validation of causal relationships on real-world datasets, **Figure 1.C**. depicts the procedure for collecting diverse attributes, scores, and tests, and **Figure 1.B**. illustrates the training process for the DNN model. **Figure 1.A**. shows the procedure for estimating causal relationships from a given data matrix. The following subsections provide a detailed explanation of each process.

**Figure 1.**
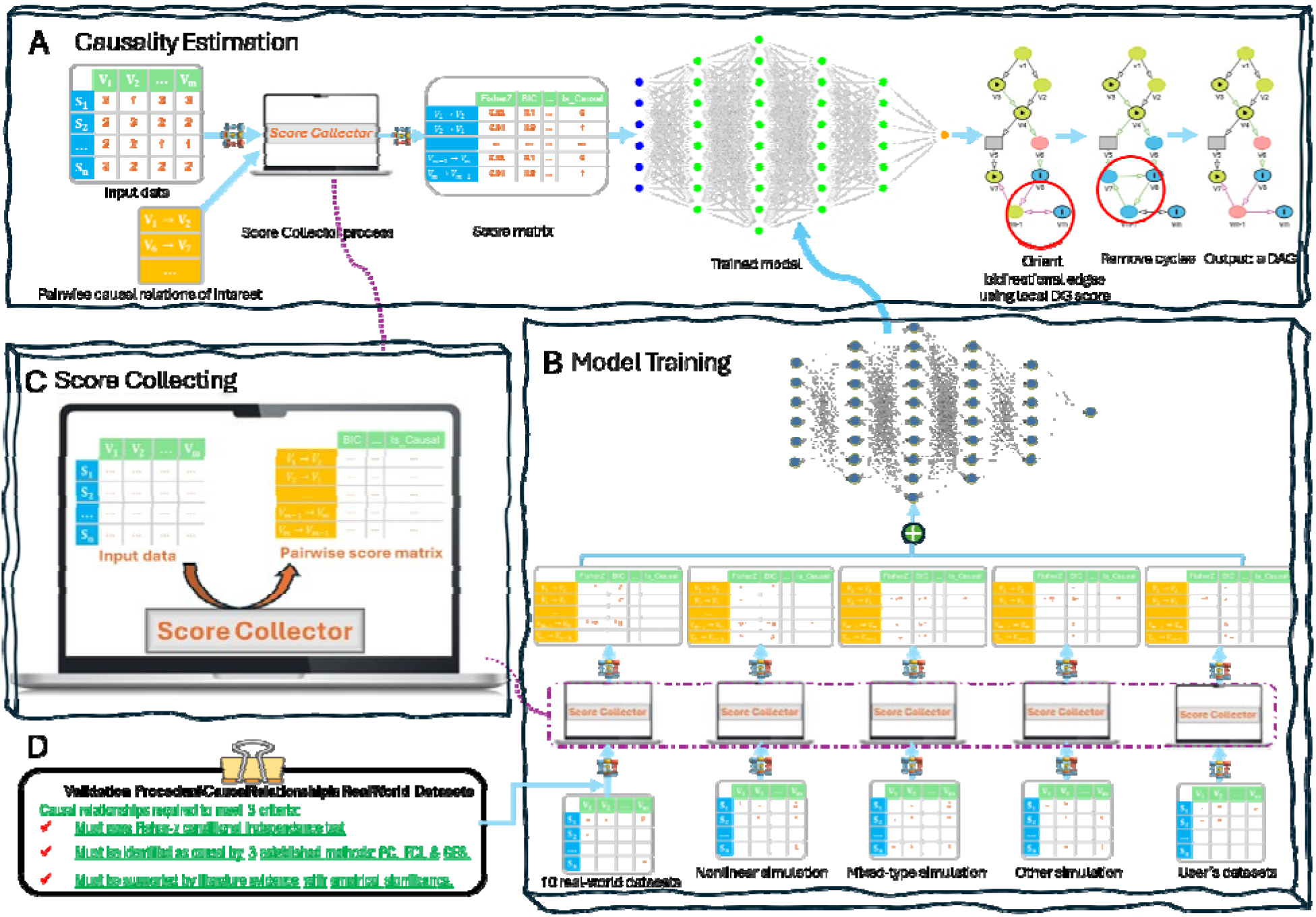
An overview of the proposed framework - DeepCEF. The figure is divided into four parts: (A) Top: Procedure for estimating causal relationships from a given data matrix; (C) Middle-Left: Collection of diverse attributes, scores, and tests; **(B)** Bottom-Right: Training process for the DNN model; and (D) Bottom-Left: Validation of causal relationships on real-world datasets.

### Data Normalization

Data normalization is a crucial preprocessing step in training DNNs due to its significant impact on the convergence, stability, and performance of the model. Neural networks, particularly those with multiple layers, are highly sensitive to the scale and distribution of input features. As demonstrated in Supplemental Figure 1.A., normalizing data before training improves model performance, as evidenced by increased prediction accuracy.

Normalizing the input data may have contributed to this performance improvement for a variety of reasons. First, without normalization, features with larger magnitudes can dominate the learning process, leading to suboptimal model performance and prolonged training times. Also, if the input features are not properly scaled, the gradients propagated through the layers during backpropagation can either vanish (become extremely small) or explode (become extremely large). This issue is exacerbated in networks with many layers, as the gradients are multiplied repeatedly through the chain rule. As shown in Supplemental Figure 1.A. and B., normalization helps mitigate this problem by ensuring that the inputs to each layer are within a consistent range, thereby promoting stable gradient flow. Furthermore, normalization can improve the generalization capability of the model by reducing the risk of overfitting. When features are on different scales, the model may assign disproportionate importance to certain features, leading to poor generalization on unseen data. By normalizing the data, the model is encouraged to learn balanced representations, which can improve its ability to generalize to new datasets.

Additionally, the activation functions used in this study, such as the sigmoid, tanh, and Rectified Linear Unit (ReLU), are sensitive to the scale of their inputs. For instance, the sigmoid and tanh functions saturate when inputs are too large or too small, resulting in gradients close to zero and hindering learning. Normalization ensures that the inputs to these activation functions remain within an optimal range, allowing them to operate effectively and maintain meaningful gradients. Finally, the initialization of weights in a neural network plays a critical role in determining the success of training. When input features are not normalized, the initial weights may interact poorly with the data, leading to unstable training dynamics. Normalization reduces the dependence on careful weight initialization, making the training process more robust and less sensitive to the choice of initial parameters.

In this study, we employed min-max normalization to rescale the features to a fixed range of [0, 1], ensuring that all features contributed equally to the learning process (as shown in Figure 1.B.). The min-max normalization formula is given by:

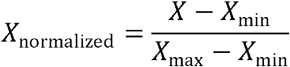

where *X* represents the original feature value, and *X*_min_ and *X*_max_ are the minimum and maximum values of the feature, respectively. This method preserves the original distribution of the data while ensuring that all features are on a comparable scale.

This preprocessing step was critical for achieving efficient training and optimal performance of the DNN. By normalizing the data, we ensured that the model could learn effectively from the input features, leading to faster convergence, improved stability, and better generalization.

### Score Collecting

Real-world observational datasets, especially those derived from complex biological systems, often encompass diverse data types and intricate relationships. This complexity necessitates the development of a more comprehensive analytical approach. To address this challenge, as illustrated in Figure 1.C., our framework integrates a broad spectrum of local causality estimation scores, independence tests, and variable attributes, enabling it to capture a wide variety of causal mechanisms for every pairwise relationship.

Our score collecting process takes as input a data matrix X ∈ R^n×d^, where *n* represents the number of observations and *d* the number of variables, along with a set of pairwise relationships *S*. The nature of the problem depends on the specification of *S*. When *S* consists of user-defined pairwise relationships or relationships associated with a target variable *R*, the task becomes a causal inference problem. In this case, the goal is to infer causality over the specified relationships in *S*. Conversely, when *S* is not predefined and must be learned from the data, the task transforms into a causal discovery problem. Here, the objective is to uncover the underlying causal structure of the system directly from the observational data *X*.

As discussed in the Data Normalization section, normalization ensures that the collected data fall within a consistent range, enhancing the model’s generalization capability on unseen data. Therefore, the input data *X* are normalized prior to the scoring process.

If two variables fail conditional independence tests, indicating the absence of a causal relationship, we classify the relationship as non-causal and exclude it from the scoring process.

We organize the collected information into three categories: (1) (conditional) independence tests, (2) context information, and (3) causal direction estimators. Each category provides unique insights, equipping the model to uncover the underlying patterns of the data generation processes.

#### (Conditional) Independence tests: (Conditional) Independence tests

are fundamental to causal inference and discovery, as they help identify relationships between variables and distinguish between mere associations and causal effects. In causal inference, these tests assess whether two variables are independent given a set of conditioning variables, which is critical for identifying and controlling for confounding factors—a key step in establishing causal relationships. In causal discovery, conditional independence tests serve two primary purposes: (1) pruning edges in a graph, where the absence of a direct causal relationship between two variables is inferred if they are conditionally independent given other variables; and (2) orienting edges, where conditional independence relationships help determine the direction of causal effects by identifying colliders and ruling out specific causal structures. However, many conditional independence tests rely on assumptions of linearity or specific functional forms, which may not hold in practice. Furthermore, violations of the faithfulness assumption can lead to incorrect causal conclusions.

In this study, we have compiled a comprehensive collection of (conditional) independence tests, including the Fisher-z test (Fisher, Ronald Aylmer 1921, 3–32), Hilbert-Schmidt Independence Criterion (HSIC) (Gretton et al. 2005, 63–77), Kernel-based Conditional Independence test (KCI) (Zhang et al. 2011, 804–813), Missing-value Fisher-z, and their variants. A complete list of the (conditional) independence tests collected in this study is provided in Supplemental Table 2.

#### Context information

Many local scoring methods excel in specific data types or application domains but struggle in others, limiting their ability to capture the complexity of underlying causal mechanisms. A method may perform well on linear relationships or continuous variables but fail to handle discrete variables or nonlinear relationships effectively. To enable the DNN model to identify these complex patterns, we have collected various contextual factors that reflect the underlying data characteristics, such as variable data types (discrete or continuous) and pairwise relationship types (linear or nonlinear). A complete list of the context factors collected in this study is provided in Supplemental Table 2. We employ the Lagrange Multiplier (LM) test (Cox and Hinkley 1979) and its variants to assess whether a pairwise relationship is nonlinear. The LM test is a hypothesis testing procedure that evaluates whether a more complex model (e.g., a nonlinear model) provides a significantly better fit to the data compared to a simpler model (e.g., a linear model). It tests the null hypothesis that the simpler model is adequate against the alternative hypothesis that the more complex model is necessary.

#### Causal direction estimators

In causal inference and discovery, various methods are utilized to assess relationships between variables, identify local causal structures, and evaluate model fit, which include negative k-fold cross-validated log likelihood (Huang et al. 2018, 1551–1560), Mutual Information (MI) (Kraskov, St ogbauer, and Grassberger 2004, 066138; Ross 2014, 1–5; Kozachenko 1987, 9), Conditional Mutual Information (CMI) (Steeg and Galstyan a; Steeg and Galstyan b), Additive Noise Model (ANM) (Hoyer et al. 2008), Bayesian Information Criterion (BIC) (Schwarz 1978, 461–464), and Degenerate Gaussian (DG) (Andrews, Ramsey, and Cooper 2019, 4–21).

Negative k-fold cross-validated log likelihood (Huang et al. 2018, 1551–1560) is used to validate causal models by measuring how well they predict outcomes under different interventions or conditions.

MI identifies potential causal relationships by detecting strong dependencies between variables. For example, a high *MI*(*X,Y*) suggests *X* and *Y* are related, though it does not imply causation. CMI tests for conditional independence, which is usually utilized in causal inference or discovery. For instance, it is used in methods like the PC algorithm (Spirtes, Peter, Glymour, and Scheines 2000) and FCI algorithm (Spirtes, Peter L., Meek, and Richardson 2013) to control for confounding and infer causal effects. If *CMI*(*X,Y*∣*Z*)=0, *X* and *Y* are conditionally independent given *Z*, which helps rule out spurious associations. ANM is used for inferring causal directions in bivariate settings by testing whether *Y*=*f*(*X*)+*∈* fits the data better than *X*=*g*(*Y*)+*∈*′, where *∈* and *∈*′ are independent noise terms. ANM also helps identify causal relationships in complex systems where the effect is a nonlinear function of the cause. BIC can be used as a criterion to compare different causal models and select the one that best explains the data without overfitting, and it is also used in structural equation modeling (SEM) to choose the best local structure among competing causal structures. The DG distribution is a special case of the multivariate Gaussian distribution where it is useful in settings where the data lies on a lower-dimensional manifold, allowing for more efficient estimation of causal effects.

By leveraging these techniques, DNN models can more accurately uncover causal relationships and understand the mechanisms underlying observed data.

### Model Selection and Training

Given a data matrix X ∈ R^n×d^, where *n* represents the number of observations and *d* the number of variables, our goal is to train a model capable of inferring causality with respect to a target variable or a set of pairwise relationships, as well as discovering the underlying causal structure. To enhance the robustness and generalizability of the final trained model, we employ various optimization techniques and integrate a diverse range of simulation data alongside 10 real-world datasets during the training process. The following steps outline our approach to obtaining the final trained model.

#### 1) Dataset Generation and Preprocessing

We generated five simulation datasets (see **Causal Data Simulation**), each comprising 1,500 observations, 50 variables, and 100 predefined causal relationships. These datasets include one generated using the Linear Fisher Model (FISHER, R. A. 1936, 179–188), three using the Mixed Graphical Model (Lee, Jason D. and and 2015, 230–253), and one nonlinear dataset. Additionally, we incorporated 10 real-world datasets with validated causal relationships (see **Validated Causal Relationships Derived from Real-World Datasets**). All datasets were rescaled using min-max normalization (see **Data Normalization**).

#### 2) Score Collection and Data Preparation

For each dataset, we collected local scores, conditional independence test results, and attributes for every pairwise relationship (see **Score Collecting**). To create a balanced training set, we introduced an equivalent number of false causal relationships into the score data, derived from the p-values of conditional independence tests. These false relationships served as negative controls for model training. Subsequently, we merged the score data across all datasets to create a unified training dataset.

#### 3) Model Selection and Evaluation

Our final model is designed as a binary classification model to accurately determine whether a pairwise relationship between two variables is causal. To identify the most suitable model, we evaluated four well-established algorithms: Support Vector Machine (SVM) (Cortes and Vapnik 1995, 273–297), Deep Neural Decision Tree (DNDT) (Yang, Morillo, and Hospedales 2018), Deep Neural Decision Tree Forest (DNDTF) (Kontschieder et al. 2015, 1467–1475), and Multilayer Perceptron (MLP). These models were tested on a simulation dataset containing linear, nonlinear, and mixed-type data. The MLP outperformed the other models in predicting true causal relationships (see **Supplemental Figure 3**) and demonstrated superior precision (see **Supplemental Figure 4**). Consequently, the MLP was selected as the default model for this study. However, to provide flexibility, the other three models were also integrated into our published software package, allowing users to choose alternative models for their analyses.

#### 4) MLP Architecture and Training

**Supplemental Figure 5** provides an overview of the MLP architecture used in this study. The network consists of an input layer, multiple hidden layers, and an output layer. The input layer accepts feature vectors of dimension *d*, where *d* corresponds to the number of input features (scores, test results, and attributes). The hidden layers, indexed by *l*, contain *n*_*l*_ neurons each and employ a variety of activation functions, including Rectified Linear Unit (ReLU) (Fukushima 1969, 322–333), Swish (SiLU) (Ramachandran, Zoph, and Le 2017), Hyperbolic Tangent (tanh), and Sigmoid. The output layer consists of two neurons, corresponding to the binary classification task (causal or non-causal). The network is trained using the backpropagation algorithm with the AdamW optimizer (Loshchilov and Hutter 2017) to minimize the binary cross-entropy loss. Dropout is applied during training to mitigate overfitting and enhance generalization performance.

#### 5) Model Implementation and Availability

After training, the network architecture, learned weights, optimizer state, and evaluation metrics are saved in a unified format for reproducibility. The model is implemented using the TensorFlow deep learning framework. The complete implementation, along with detailed documentation, is available at the GitHub repository: https://github.com/ZhenjiangFan/DeepCEF.

### Causality Estimating

As illustrated in **Figure 1.A**., estimating causal relationships from an observational dataset involves five key steps: (1) normalizing the input data, (2) collecting score information (scores, tests, and attributes), (3) loading and running the model on the score data, (4) creating a directed graph and orienting bidirectional edges, and (5) removing cycles to output a DAG. **Step 1: Data Normalization**. The process begins with an input data matrix X ∈ R^n×d^, where *n* represents the number of observations and *d* the number of variables, along with a set of pairwise relationships *S*. The set *S* can be derived from three scenarios:

1. **Targeted Causal Inference:** When a target variable *R* is specified, *S* consists of pairwise relationships associated with *R*.
2. **Focused Causal Inference:** When the user is interested in a specific set of relationships, *S* is defined as that set.
3. **Causal Discovery:** When no specific relationships are targeted, *S* includes all possible pairwise relationships.

The first two scenarios are categorized as **causal inference** tasks, while the third is regarded as **causal discovery**. The input data X is rescaled using the min-max normalization algorithm to ensure consistent scaling across variables.

**Step 2: Score Data Collection**. Next, score data (scores, conditional independence tests, and attributes) are collected for each relationship in S*S*. Relationships that fail to pass conditional independence tests are filtered out, ensuring that only statistically plausible relationships are retained for further analysis.

**Step 3: Model Execution**. The collected score data is then used as input to the causal model. For relationships estimated as causal, directed edges are added to a graph. This step leverages the model’s ability to distinguish causal from non-causal relationships based on the provided scores and tests.

**Step 4: Orienting Bidirectional Edges**. Bidirectional edges in the graph are oriented using a scoring mechanism. For example, given a bidirectional edge *A*↔*B*, the two possible causal relationships *A*→*B* and *A*←*B* are evaluated. The relationship with the higher score is selected as causal, as it is theoretically more likely to represent the true causal direction, while the other is discarded.

**Step 5: Cycle Removal and DAG Output**. Finally, cycles in the directed graph are identified and removed using a graph search algorithm. If a cycle is detected in the directed graph, the edge with the lowest **DG score** is removed to break the cycle and ensure the graph remains acyclic. This ensures that the output is a valid DAG, as shown in **Figure 1.A**., which represents the estimated causal structure of the data.

### 2.2 Causal Data Simulation

While real-world datasets are invaluable for training DNNs, they often come with challenges such as data imbalance, noise, missing annotations, and biases. To enhance the robustness and generalizability of the framework, we integrate a wide variety of simulation data. This includes data generated from the Linear Fisher model (FISHER, R. A. 1936, 179–188), the Lee and Hastie Mixed Graphical model (FISHER, R. A. 1936, 179–188; Lee, Jason D. and Hastie 2015, 230–253), and multiple nonlinear functions.

#### Simulating Causal Data Using Linear Fisher Model

We use a linear regression model from Ronald A. Fisher’s framework in this simulation scenario. The model is described as the following:

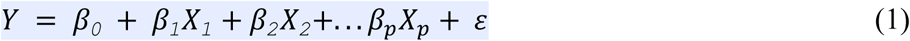

where *Y* is the dependent variable, *X*_*1*_, *X*_*2*_, …, *X*_*p*_ are the independent variables, *β*_*0*_ is the intercept, *β*_*1*_, *β*_*2*_, …, *β*_*p*_ are the slopes (regression coefficients) representing the change in *Y* for a unit change in *X*_*i*_, and *ε* is the error term (assumed to be normally distributed with mean 0 and constant variance *σ*^2^).

This project uses a Linear Fisher model implementation from Tetrad (https://www.cmu.edu/dietrich/philosophy/tetrad/), a software suite for causal model analysis. Simulations from the Linear Fisher model all feature 50 variables across 1500 samples. All parameters are set to Tetrad’s defaults for our use.

### Simulating Causal Data Using Mixed Graphical Model

Mixed Graphical Model (MGM) was proposed by Lee and Hastie (FISHER, R. A. 1936, 179– 188; Lee, Jason D. and Hastie 2015, 230–253) to learn the structure of a pairwise graphical model over continuous and discrete variables. The structure used to represent the simulated causal graph contains three relation types: 1) causal relationships between two continuous variables which are based on a multivariate Gaussian (MG) model, 2) causal relationships between two discrete variables which are based on the usual discrete pairwise Markov random field (MRF) model, 3) and mixed causal relationships between a continuous variable and a discrete variable. The mixed causal relationships have the desirable property which has the two types of conditional distributions: simple Gaussian linear regressions and multiclass logistic regressions.

For all the relations among discrete variables, the joint probability distribution of the MRF is given by the Gibbs distribution:

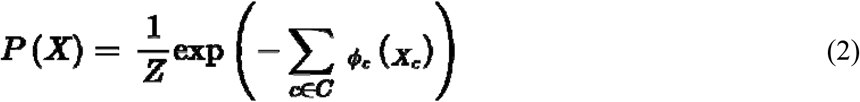

where *X* = (*X*_1_, *X*_2_, …, *X*_*n*_) is the set of all random discrete variables, *C* is the set of the cliques around all the relations (second-order), *ϕ*_*c*_(*X*_*c*_) is the potential function for clique *c*, and *Z* is the partition function (a normalization constant ensuring the distribution sums to 1).

The implementation of this MGM simulation is also from Tetrad, where each simulation dataset has a shape of 50 variables and 1500 samples. We use the default values for all the simulation parameters.

### Simulating Nonlinear Causal Data Using Trigonometric, Hyperbolic, and Power Functions

Nonlinear relationships are fundamental to modeling complex systems across various scientific and engineering domains. To generate synthetic nonlinear data that captures a wide range of behaviors, we employ a combination of trigonometric, hyperbolic, and power functions. These functions are chosen for their ability to model diverse nonlinear patterns, including periodic oscillations, sigmoidal transitions, power-law relationships, and bounded responses. Below, we describe the role of each function and how they can be combined to simulate complex nonlinear data.

#### Arctangent Function (arctan)

The arctan function, or inverse tangent, produces a smooth, S-shaped curve that saturates as the input approaches infinity. It is bounded between 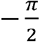 and 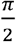, making it ideal for modeling saturating behaviors. In this work, this function was scaled and shifted to fit specific data ranges:

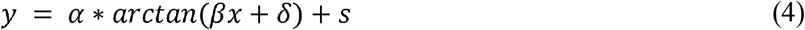

where *α, β, δ*, and *s* are parameters controlling amplitude, steepness, horizontal shift, and vertical shift, respectively.

#### Cosine Function (cos)

The cos function introduces periodic oscillations into the data. It is useful for simulating cyclical patterns, such as seasonal trends or wave-like behavior. By adjusting the frequency, amplitude, and phase, you can control the period and magnitude of the oscillations:

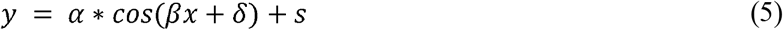

where *α* controls amplitude, *β* controls frequency, *δ* controls phase shift, and *s* controls vertical shift.

#### Arcsine Function (arcsin)

The arcsin function, or inverse sine, produces a nonlinear relationship bounded between 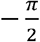 and 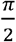. It is useful for modeling data with a restricted range. To generalize, we scale and shift the function:

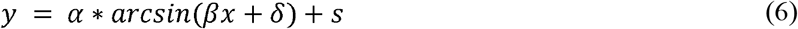

where *α, β, δ*, and *s* are parameters controlling amplitude, steepness, horizontal shift, and vertical shift, respectively.

#### Power Function (power)

The power function, *y* = *x*^*n*^, allows for the creation of polynomial-like relationships. By varying the exponent *n*, you can control the degree of nonlinearity. This function is useful for modeling accelerating or decelerating trends.

#### Sine Function (sin)

The sin function, like cos, introduces periodic behavior but with a phase shift. It is often used in combination with cos to model more complex waveforms. We adjust its parameters similarly to the cos function:

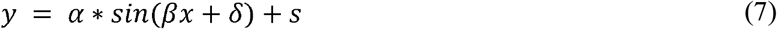

where *α* controls amplitude, *β* controls frequency, *δ* controls phase shift, and s controls vertical shift.

#### Hyperbolic Tangent Function (tanh)

The tanh function is a smooth, S-shaped function bounded between −1 and 1. It is commonly used in machine learning and data modeling due to its saturating properties. This function is particularly useful for simulating data that plateaus at extreme values. It can be scaled and shifted as follows:

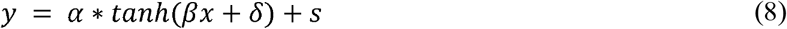

where *α, β, δ*, and *s* are parameters controlling amplitude, steepness, horizontal shift, and vertical shift, respectively.

#### Arccosine Function (arccos)

The arccos function, or inverse cosine, produces a nonlinear relationship bounded between 0 and π. It is useful for modeling data with a restricted range. To generalize, we scale and shift the function:

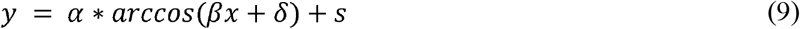

where *α, β, δ*, and *s* are parameters controlling amplitude, steepness, horizontal shift, and vertical shift, respectively.

#### Combining Functions for Complex Nonlinearity

These functions can be combined to simulate more complex nonlinear data. This combination introduces saturating, periodic, and smooth transition behaviors into the data. Additionally, scaling and shifting parameters can be incorporated to fine-tune the simulation:

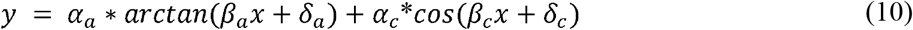

where *α*_*a*_, *β*_*a*_, *δ*_*a*_, *α*_*c*_, *β*_*c*_, and *δ*_*c*_ are adjustable parameters.

The data generated using these functions can be used to test the robustness of machine learning models, validate causal inference algorithms, or explore the behavior of nonlinear systems in controlled settings. Supplemental Figure.1 depicts the nonlinear trend of the simulation data generated from these functions. The applications for such a simulation approach include: 1) simulating biological systems with oscillatory dynamics and saturation effects; 2) modeling physical systems with power-law scaling and bounded responses; and 3) testing the performance of regression models on data with diverse nonlinear patterns.

By leveraging the unique properties of these functions, we can create realistic and diverse nonlinear datasets that mimic the complexities of real-world phenomena. This approach provides a flexible framework for generating synthetic data tailored to specific research needs, enabling rigorous testing and validation of analytical methods.

### 2.3 Validated Causal Relationships Derived from Real-World Datasets

To enhance the robustness and generalizability of the framework, we integrate a wide variety of simulation data. This includes data generated from the Linear Fisher model, the Lee and Hastie Mixed Graphical model, and multiple nonlinear functions.

The efficacy of DNNs is heavily dependent on the quality, diversity, and scale of the datasets used for training. Real-world datasets play a pivotal role in enabling DNNs to generalize well to unseen data, as they capture the inherent variability, noise, and complexity of practical scenarios. This section introduces 10 prominent real-world datasets (Supplemental Table 1) and some of them have been widely used to train and evaluate deep neural in the field computational biology.

Supplemental Table 1 details the 10 real-world biological datasets used in the training process, including disease type, data type, data source, access links, and other relevant information. Among these datasets, six consist of clinical and laboratory data, one comprises long-read sequencing data, and three are single-cell sequencing datasets. For clinical and laboratory datasets with missing values, we employed imputation strategies such as mean imputation and K-Nearest Neighbors. Additional details on the preprocessing steps are available in our GitHub repository.

The causal relationships incorporated into the training procedure were required to meet three criteria:

1. They must pass the Fisher-z conditional independence test to demonstrate a statistical association.
2. They must be identified as causal by three established causal discovery methods: the PC algorithm (FISHER, R. A. 1936, 179–188; Lee, Jason D. and Hastie 2015, 230–253; Spirtes, Peter, Glymour, and Scheines 2000), FCI (FISHER, R. A. 1936, 179–188; Lee, Jason D. and Hastie 2015, 230–253; Spirtes, Peter L., Meek, and Richardson 2013), and GES (FISHER, R. A. 1936, 179–188; Lee, Jason D. and Hastie 2015, 230–253; Spirtes, Peter L., Meek, and Richardson 2013; Chickering 2002, 507–554).
3. They must be supported by literature evidence confirming a significant relationship between the variables.

For each relationship satisfying the first two criteria, we conducted a literature search using Scale AI’s tool (for reproducibility) and Google Scholar to identify relevant publications that study the relationship. Supplemental Material 1 provides a comprehensive list of these relationships, along with details of the supporting studies. The directed causal relationships for each dataset are illustrated in Supplemental Material 2.

## Results

### 3.1 Simulation Results

To evaluate the effectiveness of DeepCEF, we conducted extensive simulations and compared its performance against three widely used methods: GES, FCI, and PC, focusing on prediction accuracy and precision. We generated 20 simulation datasets (see **Causal Data Simulation**), each comprising 1,500 samples and 90 variables. Each dataset included 60 linear, 60 nonlinear, and 60 mixed-type true causal relationships. The implementations of GES, FCI, and PC were obtained from the causality package Tetrad (https://github.com/cmu-phil/tetrad).

As shown in **Figure 2.A**., our method consistently achieved superior accuracy across all 20 simulation datasets, with a prediction accuracy rate consistently above 90%. In contrast, GES, FCI, and PC achieved accuracy rates of approximately 63%, 50%, and 69%, respectively. All methods exhibited consistent performance, with small variations in accuracy rates across datasets. Notably, PC outperformed GES and FCI, which may be attributed to the use of the latest version of the PC algorithm from Tetrad.

**Figure 2.**
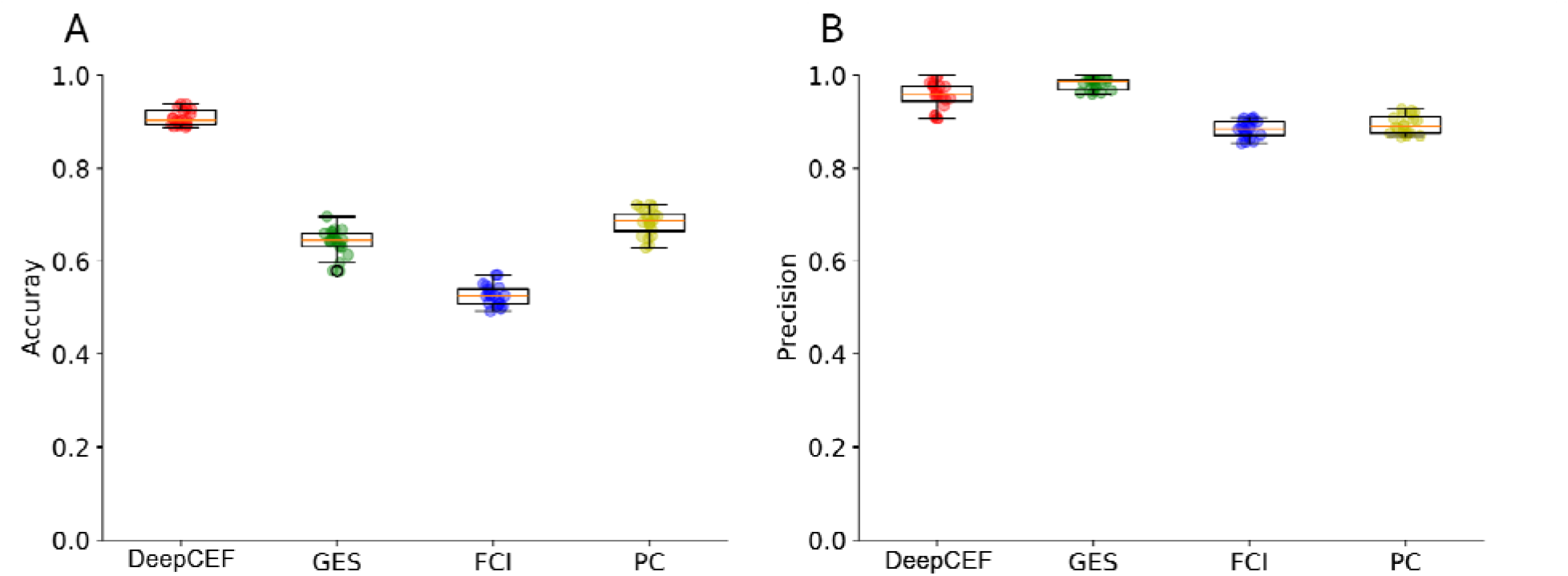
Performance comparison between DeepCEF and existing methods (GES, FCI, and PC) in terms of prediction accuracy and precision. (A) DeepCEF outperforms GES, FCI, and PC in predicting the number of true causal relationships across 20 simulation datasets. (B) DeepCEF achieves prediction precision comparable to GES and higher than FCI and PC across 20 simulation datasets.

In terms of prediction precision as shown in Figure 2.B., GES achieved relatively high precision (around 98%), while our method demonstrated comparable precision across all datasets. In contrast, FCI and PC performed poorly in this regard.

In summary, the simulation studies demonstrate that DeepCEF significantly outperforms existing methods (GES, FCI, and PC) in terms of accuracy, even in complex and noisy settings. This robust performance highlights the potential of our method for practical applications in causal inference and discovery.

### 3.2 Real-world Application Results

#### PAM50

To evaluate the performance of DeepCEF in learning causal relationships from complex real-world data, we compared it with three established methods—GES, FCI, and PC—using a breast invasive carcinoma (BRCA) dataset from Fan et al ((Fan et al. 2023, gad044)). The dataset, originally sourced from The Cancer Genome Atlas (TCGA), is available through the UCSC Xena browser. The preprocessed data used in this study includes 601 samples and 503 features, comprising 502 genes and one clinical variable, PAM50. PAM50 (Parker et al. 2009, 1160–1167) is a critical clinical feature for categorizing breast tumors based on the expression of 50 predefined genes (PAM50 genes). We consider causal directions from these genes to PAM50 status as true causal relationships. The evaluation metric is the number of PAM50 genes identified as having a causal relationship with PAM50.

As shown in **Figure 3.A**, DeepCEF method identifies 11 genes causally related to PAM50, significantly outperforming GES, FCI, and PC, which identify only 7, 2, and 1, respectively. This demonstrates the DeepCEF’s superior ability to uncover known causal relationships in complex datasets.

**Figure 3.**
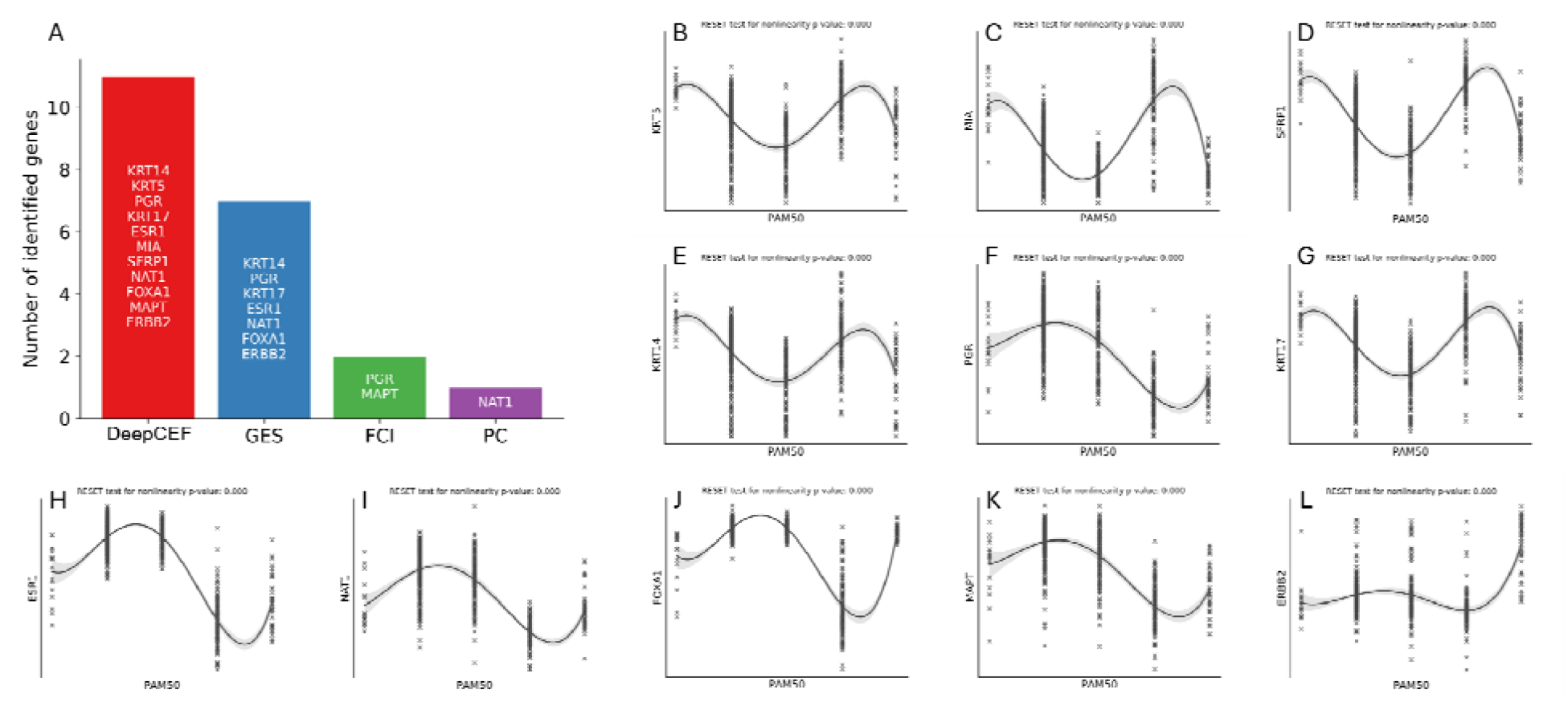
Performance comparison between DeepCEF and existing methods (GES, FCI, and PC) in identifying causal relationships and assessing nonlinear trends between PAM50-related genes and PAM50 in a BRCA dataset. (A) DeepCEF identifies 11 genes causally related to PAM50, while GES, FCI, and PC identify only 7, 2, and 1, respectively. (B–L) Nonlinear causal relationships between PAM50 and the 11 identified genes (KRT5, MIA, SFRP1, KRT14, PGR, KRT17, ESR1, NAT1, FOXA1, MAPT, and ERBB2). These nonlinear relationships explain why existing methods struggle to detect these causal links.

To investigate why the existing methods struggle to identify these relationships, we analyzed the relationships between PAM50 and the 11 genes identified by DeepCEF. **Figure 3.B–L** reveals strong nonlinear trends between PAM50 and the identified genes (KRT5, MIA, SFRP1, KRT14, PGR, KRT17, ESR1, NAT1, FOXA1, MAPT, and ERBB2). We further validated these nonlinear relationships using Ramsey’s RESET test for nonlinearity. All tests yielded a p-value of 0, providing strong statistical evidence of nonlinearity. These results explain the limitations of existing methods, which are less effective at detecting nonlinear relationships, and highlight DeepCEF’s capability to identify both linear and nonlinear causal relationships in complex datasets.

#### Breast Cancer

To demonstrate the practical utility of DeepCEF and its ability to uncover underlying causal relationships in complex real-world systems, we applied it to a clinical breast cancer dataset. The goal was to evaluate whether the method could identify relevant causal relationships associated with breast cancer diagnosis. The dataset comprises 10 clinical variables collected from 64 breast cancer patients and 52 healthy controls. **Table 1** provides a detailed description of the variables, including their names, data types, descriptions, and units of measurement.

**Table 1.** Description of variables in the breast cancer dataset used in this study, including variable names, data types, descriptions, and units of measurement.

| Variable Name | Type | Description | Units |
| --- | --- | --- | --- |
| Age | Integer | Age | Year |
| BMI | Continuous | Body mass index | kg/m<sup>2</sup> |
| Glucose | Integer | Serum Glucose level | mg/dL |
| Insulin | Continuous | Insulin level | μU/mL |
| HOMA | Continuous | Homeostasis Model Assessment |  |
| Leptin | Continuous | Leptin level | ng/mL |
| Adiponectin | Continuous | Adiponectin level | μU/mL |
| Resistin | Continuous | Resistin level | ng/mL |
| MCP-1 | Continuous | Chemokine Monocyte Chemoattractant Protein 1 | pg/dL |
| Diagnosis of breast cancer | Integer | 1=Healthy controls, 2=Patients |  |

**Figure 4** presents the DAG generated by DeepCEF, illustrating causal relationships among clinical variables and breast cancer diagnosis.

**Figure 4.**
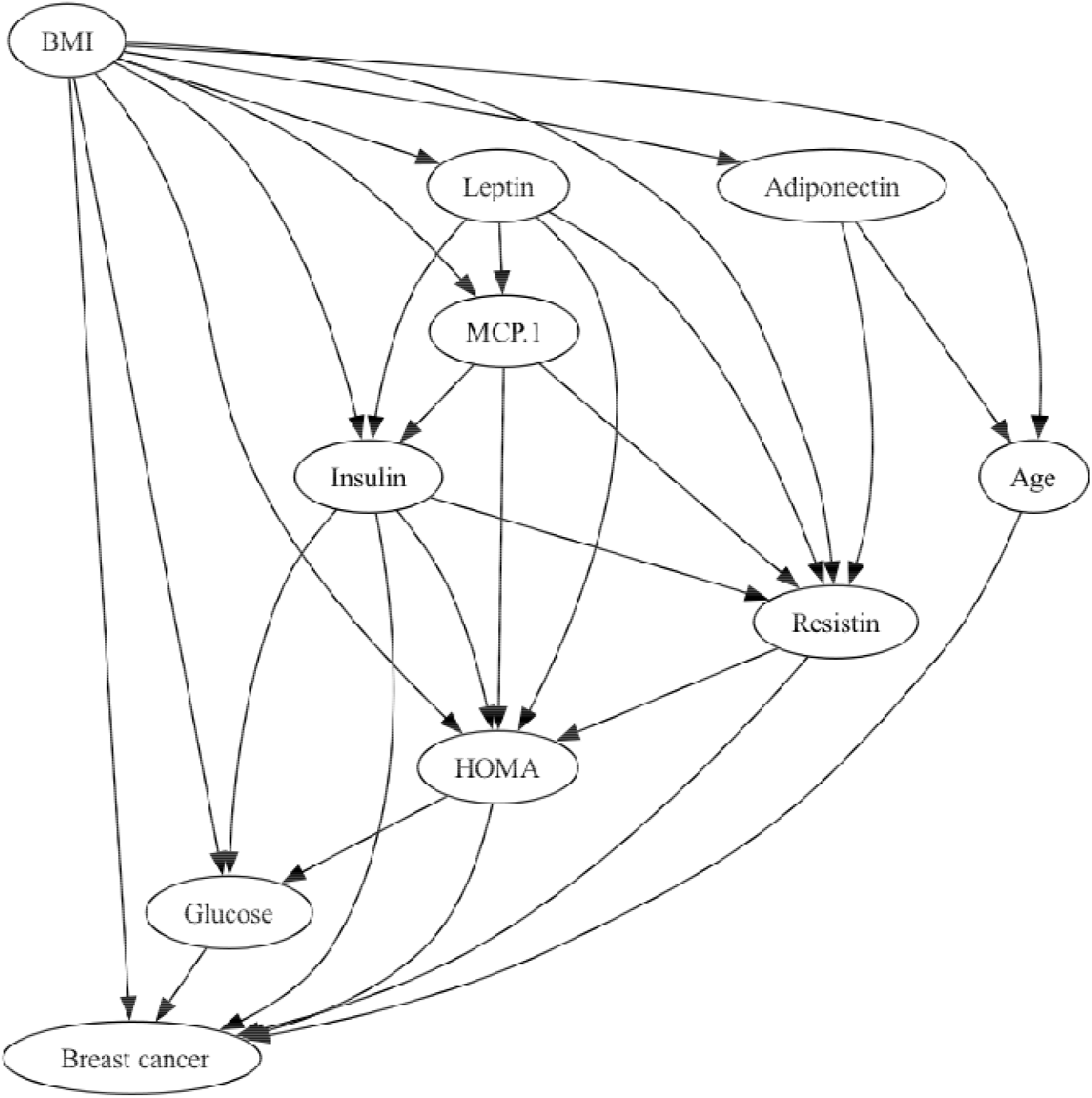
Directed acyclic graph (DAG) generated by DeepCEF, depicting causal relationships among clinical variables and breast cancer diagnosis. Six clinical variables—BMI, glucose, insulin, HOMA, resistin, and age—were identified as having causal relationships with breast cancer diagnosis. The DAG also captures other well-established causal relationships, such as the influence of insulin on glucose levels.

Six clinical variables—BMI, glucose, insulin, HOMA, resistin, and age—were identified as having causal relationships with breast cancer diagnosis. Among these, BMI, insulin, and HOMA are particularly notable due to their established links to breast cancer through mechanisms involving hormonal changes, inflammation, and metabolic dysregulation. Elevated BMI, a known risk factor for postmenopausal breast cancer, increases estrogen production by adipose tissue, promoting hormone receptor-positive cancers (Cheraghi, Zahra AND Poorolajal, Jalal AND Hashem, Tahereh AND Esmailnasab, Nader AND Doosti Irani,Amin 2012, 1–9; Guo et al. 2017, 1814–1822). Elevated insulin levels, also linked to increased breast cancer risk (especially in postmenopausal women), promote cell proliferation and inhibit apoptosis (Lawlor et al. 2008, 1133–1163; Gunter et al. 2009a, 48–60). As a measure of insulin resistance, higher HOMA values are associated with increased breast cancer risk, particularly in postmenopausal women. Insulin resistance is a key factor in breast cancer development (and 2008, 63–70; Gunter et al. 2009a, 48–60).

To validate the causal relationships identified by DeepCEF, we conducted a literature review. **Supplemental Table 3** summarizes the literature studies supporting the causal relationships between breast cancer and the six clinical variables: BMI, glucose, insulin, HOMA, resistin, and age. The DAG in **Figure 4** also captures other well-established causal relationships, such as the influence of insulin on glucose levels.

Understanding the roles of colliders, mediators, and confounders is essential for accurately interpreting the relationships between clinical variables and breast cancer diagnosis.

**Supplemental Table 4** provides a summary of the colliders, mediators, and confounders identified in the DAG produced by DeepCEF for the breast cancer dataset.

## Discussion

Our proposed causal discovery framework, DeepCEF, demonstrates superior performance compared to three widely used approaches—GES, FCI, and PC—in both simulated and real-world datasets. In simulations, our method achieves significantly higher accuracy in recovering ground-truth causal structures (Figure 2). When applied to complex BRCA data containing assumedly true causal directions around the PAM50 target variable, our method outperforms existing approaches in identifying verified causal relationships (Figure 3). Furthermore, application to a clinical breast cancer dataset revealed six clinically relevant variables with causal associations to breast cancer diagnosis. Notably, BMI, insulin, and HOMA—factors with well-documented mechanistic links to breast cancer via hormonal, inflammatory, and metabolic pathways—were correctly identified (Supplemental Table 3). These results underscore our method’s ability to uncover biologically plausible causal relationships in real-world settings, particularly for nonlinear interactions (Figure 3).

### Advantages Over Traditional Methods

Unlike constraint-based methods (e.g., PC, FCI), our approach does not require the faithfulness assumption, making it robust to violations arising from parameter cancellations or deterministic relationships. This flexibility addresses a key limitation of conventional causal discovery techniques. By leveraging DNNs, our framework adapts to complex data distributions while maintaining theoretical rigor. The integration of diverse simulated and real-world biological data during training further ensures generalizability across domains.

### Practical Utility and Extensibility

Our method provides a user-friendly, extensible platform for causal analysis in biological research, with applications ranging from gene regulatory network inference to drug discovery. The modular design allows seamless incorporation of additional validation metrics, tests, or attributes as needed. Users can also expand the training corpus with additional simulated and real-world datasets to improve robustness.

### Limitations and Future Directions

While our framework shows significant improvements over existing methods, computational efficiency remains a challenge for very large datasets due to the score collecting step even after we optimized the score collection process through parallelization techniques. One future work could incorporate more powerful AI architectures (e.g., large language models) to enhance scalability.

## Conclusion

We have presented a novel deep learning-based framework for causal inference and discovery that addresses key limitations of traditional methods. By leveraging the representational power of DNNs, our approach: (1) eliminates the need for restrictive assumptions like faithfulness that constrain conventional constraint-based methods; (2) effectively captures complex, nonlinear causal relationships; and (3) demonstrates robust performance across both simulated datasets and challenging real-world biological applications.

The framework’s strong empirical performance - validated through comprehensive benchmarking against established methods (GES, FCI, PC) and application to BRCA and clinical breast cancer datasets - highlights its ability to uncover biologically plausible causal mechanisms. Notably, our method successfully identified known risk factors (BMI, insulin, HOMA) and their causal relationships to breast cancer outcomes, demonstrating its practical utility in biomedical research. As an extensible and user-friendly platform, our framework opens new possibilities for causal analysis in diverse domains, from molecular biology (gene regulation, disease pathways) to therapeutic development (drug discovery, treatment optimization). The framework’s flexibility and performance suggest broad applicability across scientific disciplines where understanding causal mechanisms is paramount.

## Supporting information

Supplemental Material 1-Diseases Exposure Outcome Publication

## Acknowledgments

The authors acknowledge the computational support provided by Google Cloud Research Credits from GCP Education Programs and AMD’s High Performance Compute Fund.

## Supplemental Figures

**Supplemental Figure 1.**
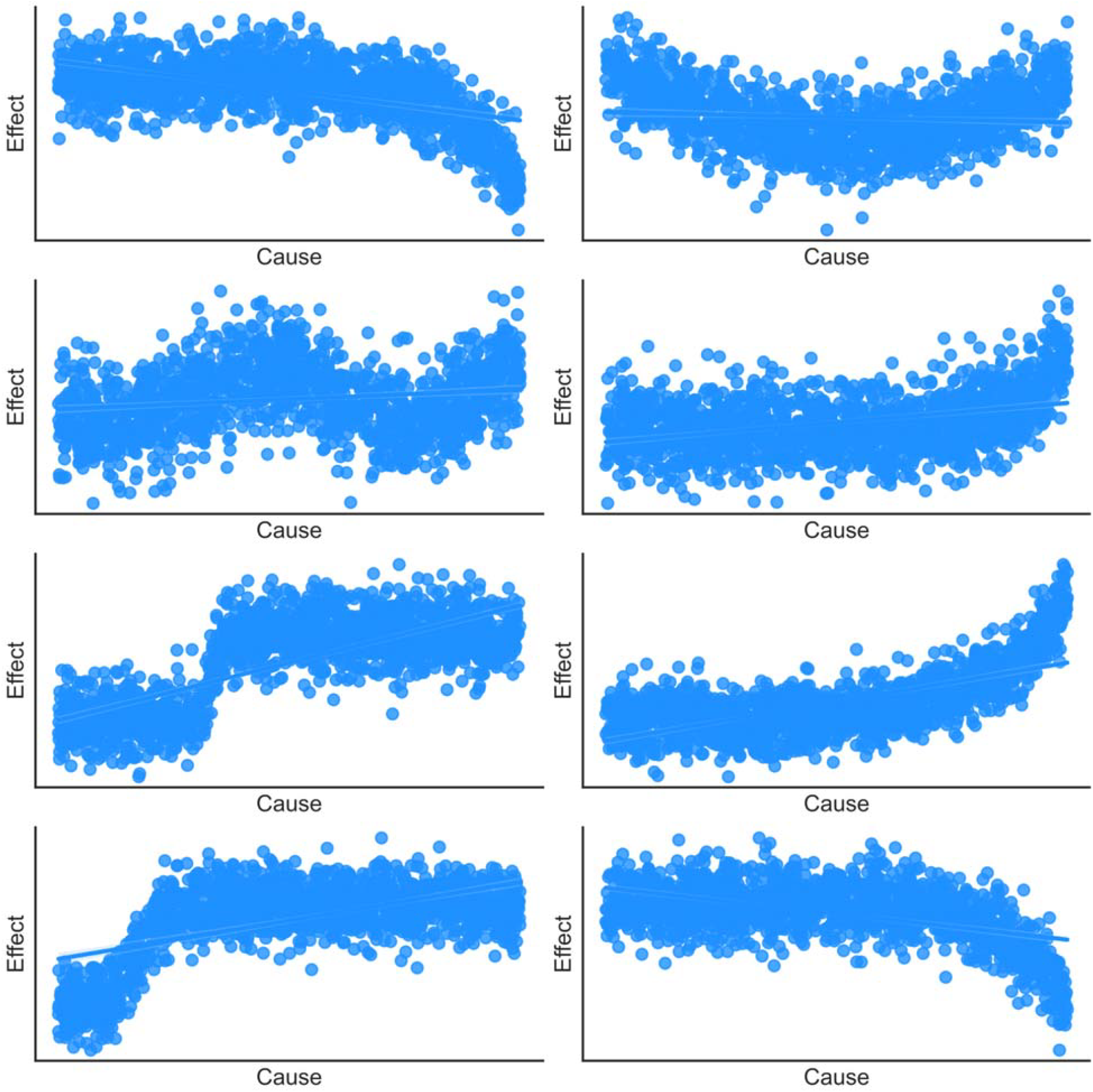
The nonlinear trend of the causal simulation data generated from nonlinear functions.

**Supplemental Figure 2.**
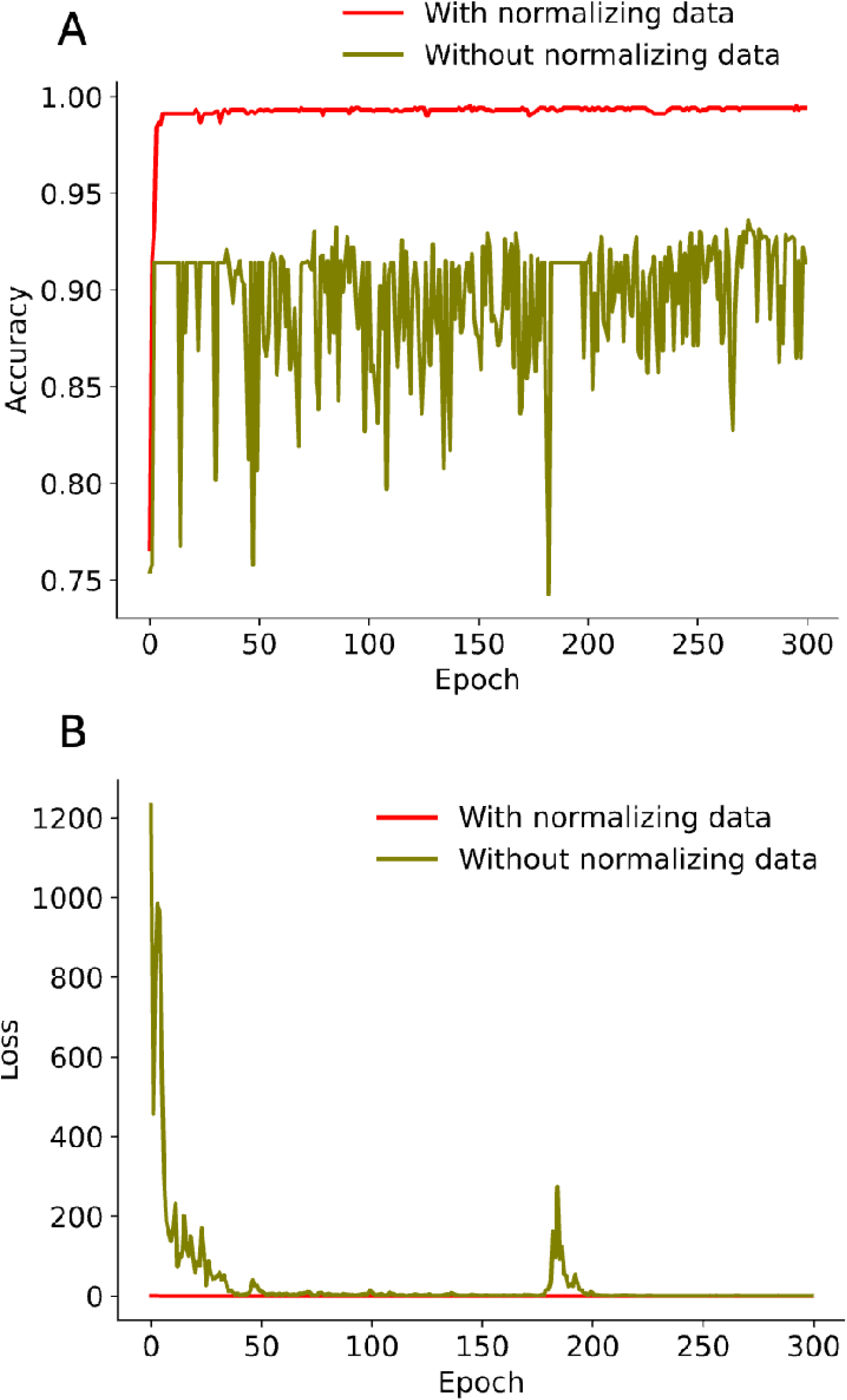
Data normalization is a critical preprocessing step for training DNNs. (A) Normalizing data prior to training enhances model performance, as measured by prediction accuracy. (B) Normalization promotes stable gradient flow during training, preventing suboptimal model performance and reducing training time.

**Supplemental Figure 3.**
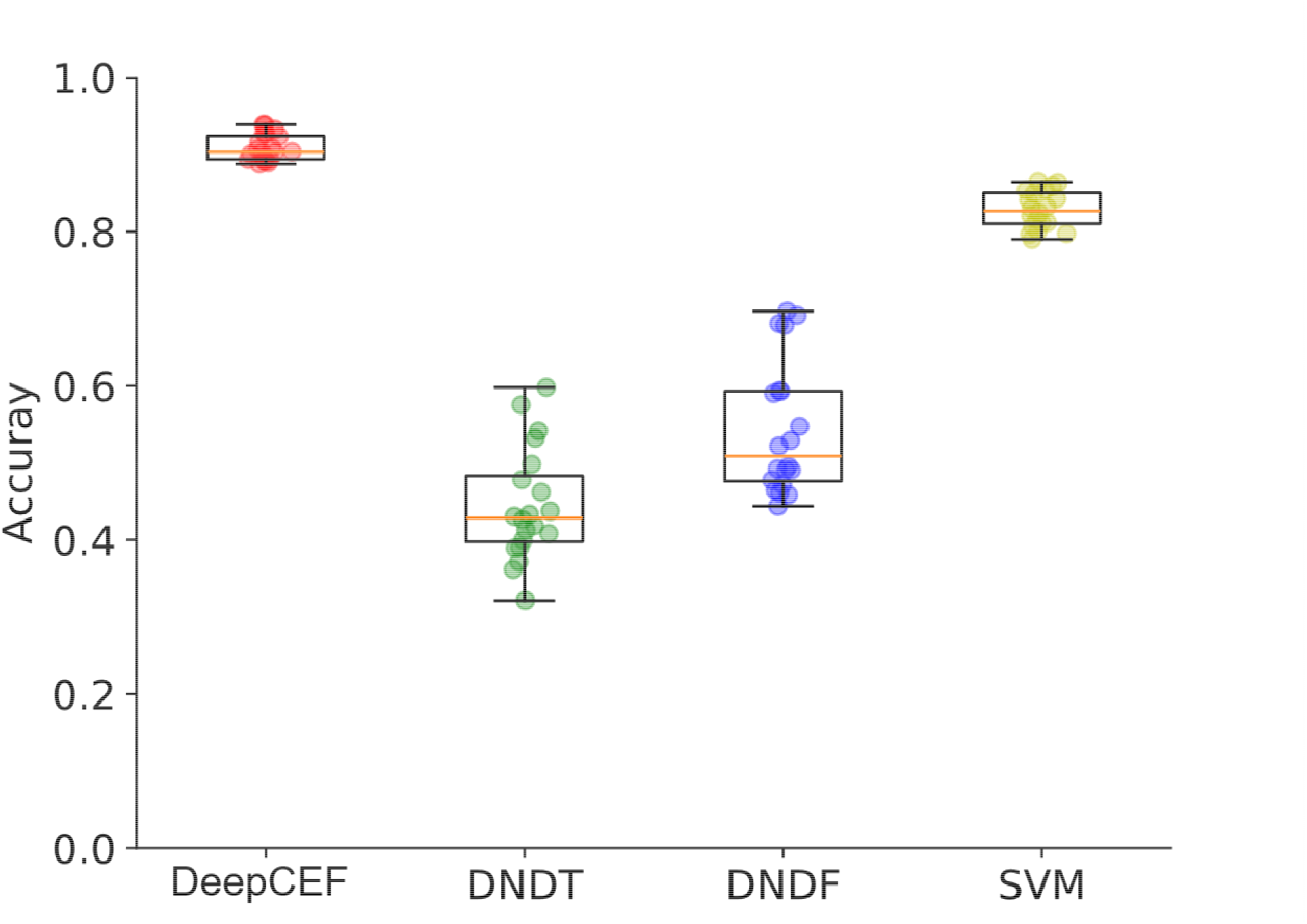
Performance comparison of the MLP against DNDT, DNDF, and SVM models in predicting the number of accurate causal relationships. The y-axis displays the distribution of 20 prediction accuracy rates, with each rate derived from a simulation dataset.

**Supplemental Figure 4.**
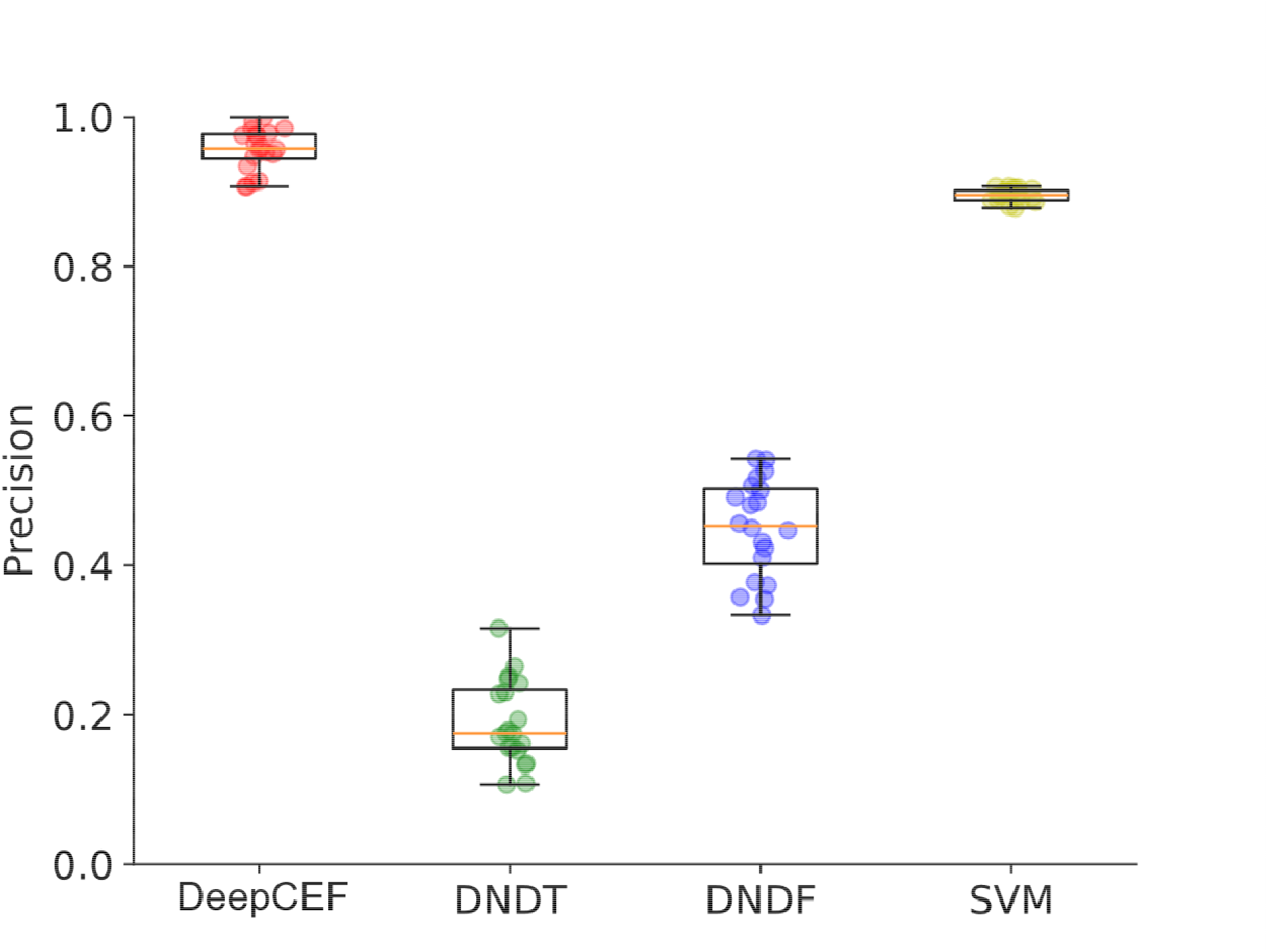
The MLP demonstrates superior prediction precision compared to DNDT, DNDF, and SVM. The y-axis represents the distribution of 20 prediction accuracy rates, each computed from a distinct simulation dataset.

**Supplemental Figure 5.**
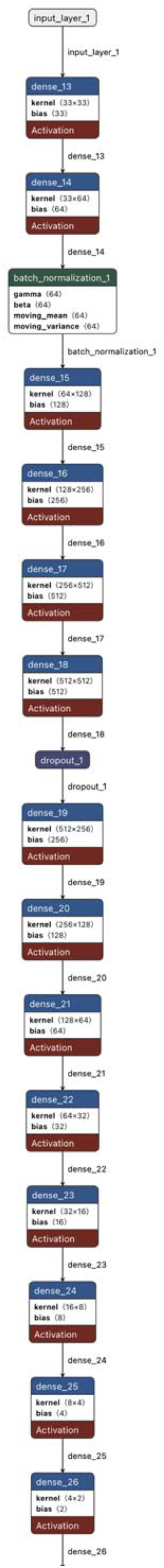
Architecture of the DNN model used in this study. The network comprises an input layer, multiple hidden layers, a dropout layer, and an output layer.

## Supplemental Tables

**Supplemental Table 1.**
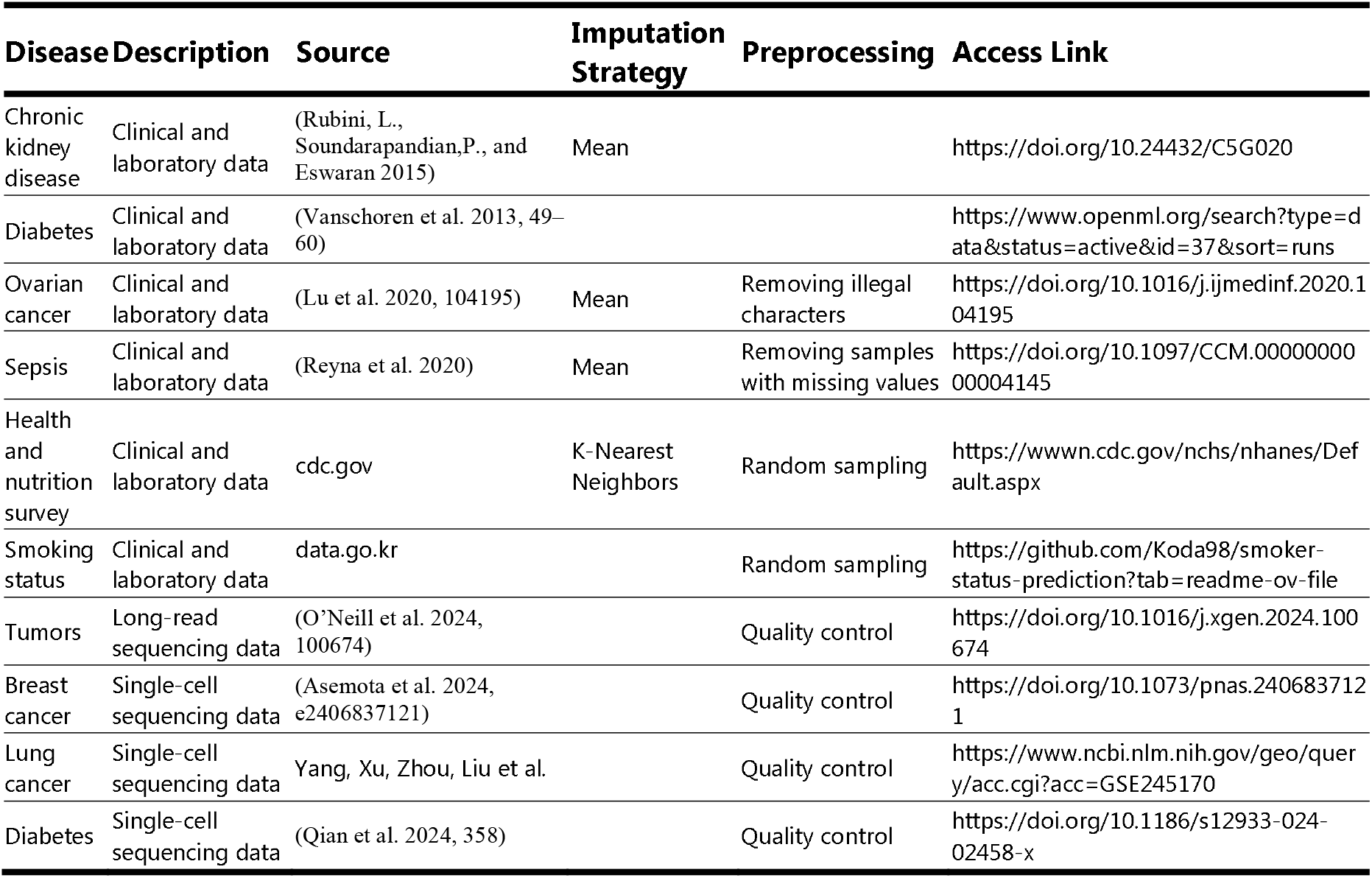
The table provides details of the 10 real-world biological datasets utilized in the training process, including disease type, data type, data source, access links, and other relevant information.

**Supplemental Table 2.** Overview of the three categories of information collected in this study: (1) (conditional) independence tests, (2) context information, and (3) causal direction estimators. The table details their applications in causal inference, associated authors or references, and their potential value ranges.

| Name | Usage | Source | Value Range |
| --- | --- | --- | --- |
| Fisher-z | Conditional independence test | (FISHER, R. A. 1936, 179–188) | 0 to 1 |
| Fisher-z (binary) | Binary label for Fisher-z conditional independence test | (FISHER, R. A. 1936, 179–188) | 0 or 1 |
| HSIC statistic | Independence test | (Gretton et al. 2005, 63–77) |  |
| HSIC p-value | HSIC Independence test | (Gretton et al. 2005, 63–77) | 0 to 1 |
| HSIC (binary) | Binary label for HSIC Independence test | (Gretton et al. 2005, 63–77) | 0 or 1 |
| KCI test | Conditional independence test | (Zhang et al. 2011, 804–813) | 0 to 1 |
| KCI (binary) | Binary label for KCI test | (Zhang et al. 2011, 804–813) | 0 or 1 |
| Missing-value Fisher-z | Conditional independence test | (FISHER, R. A. 1936, 179–188) | 0 to 1 |
| Missing-value Fisher-z (binary) | Binary label for Missing-value Fisher-z test | (FISHER, R. A. 1936, 179–188) | 0 or 1 |
| Negative k-fold cross-validated log likelihood (with regularization parameter of 0.01) | Local score | (Huang et al. 2018, 1551–1560) |  |
| Negative k-fold cross-validated log likelihood (with regularization parameter of 0.05) | Local score | (Huang et al. 2018, 1551–1560) |  |
| Negative k-fold cross-validated log likelihood (with regularization parameter of 0.1) | Local score | (Huang et al. 2018, 1551–1560) |  |
| Negative k-fold cross-validated log likelihood (binary) | Local score | (Huang et al. 2018, 1551–1560) | 0 or 1 |
| Cause variable data type (discrete or not) | Variable attribute |  | 0 or 1 |
| Cause variable data type (discrete or not) | Variable attribute |  | 0 or 1 |
| MI | Mutual information between cause and effect | (Kraskov, St ogbauer, and Grassberger 2004, 066138) |  |
| CMI | Conditional mutual information between cause and effect | (Steeg and Galstyan b) |  |
| ANM | Estimating cause and effect between two variables | (Hoyer et al. 2008) | 0 to 1 |
| LM test | Estimating linearity between two variables | statsmodels.org | 0 or 1 |
| LM test (p-value) | Determining linearity between two variables | statsmodels.org | 0 to 1 |
| LM test (p-value, augmentation type 'fitted', F-test) | Determining linearity between two variables | statsmodels.org | 0 to 1 |
| LM test (p-value, augmentation type 'exog', F-test) | Determining linearity between two variables | statsmodels.org | 0 to 1 |
| LM test (p-value, augmentation type 'princomp', F-test) | Determining linearity between two variables | statsmodels.org | 0 to 1 |
| LM test (p-value, augmentation type 'fitted'', chi-square test) | Determining linearity between two variables | statsmodels.org | 0 to 1 |
| LM test (p-value, augmentation type 'exog'', chi-square test) | Determining linearity between two variables | statsmodels.org | 0 to 1 |
| LM test (p-value, augmentation type 'princomp'', chi-square test) | Determining linearity between two variables | statsmodels.org | 0 to 1 |
| BIC (lambda 0.1) | Determining causal direction for a relation | (Schwarz 1978, 461–464) |  |
| BIC (lambda 0.5) | Determining causal direction for a relation | (Schwarz 1978, 461–464) |  |
| BIC (lambda 1) | Determining causal direction for a relation | (Schwarz 1978, 461–464) |  |
| BIC (lambda 1.5) | Determining causal direction for a relation | (Schwarz 1978, 461–464) |  |
| BIC (binary, lambda 1) | Determining causal direction for a relation | (Schwarz 1978, 461–464) | 0 or 1 |
| DG | Determining causal direction for a relation | (Andrews, Ramsey, and Cooper 2019, 4–21) |  |
| DG (binary) | Determining causal direction for a relation | (Andrews, Ramsey, and Cooper 2019, 4–21) | 0 or 1 |

**Supplemental Table 3.** Summary of literature studies supporting the causal relationships identified by DeepCEF between breast cancer and six clinical variables: BMI, glucose, insulin, HOMA, resistin, and age.

| **Biomarkers** | **Publications** |
| --- | --- |
| Glucose | (Muti et al. 2002, 1361–1368)<br>(Minicozzi et al. 2013, 3881–3888)<br>(Hiba Mahgoub and Areeg 2022)<br>(Shin and Koo 2021)<br>(Monzavi-Karbassi et al. 2016, 7)<br>(Li et al. 2024)<br>(Barba et al. 2012, 1838–1845)<br>(Luque et al. 2017, 81462–81474)<br>(Mink et al. 2002, 349–352) |
| BMI | (Smith et al. 2021, 215–223)<br>(Gallagher and LeRoith 2015, 727–748)<br>(Roberts, Dive, and Renehan 2010, 301–316)<br>(Picon-Ruiz et al. 2017, 378–397)<br>(Kolb and Zhang 2020)<br>(Engin 2017, 571–606)<br>(Nguyen et al. 2023, 4418)<br>(Dehesh et al. 2023, 392)<br>(Mohanty and Mohanty 2021, 117–123)<br>(García-Estévez et al. 2021)<br>(Blair et al. 2019, 33)<br>(James et al. 2015, 705–720) |
| Age | (Benz 2008, 65–74)<br>(Hendrick et al. 2021, 4384–4392)<br>(Giannakeas, Lim, and Narod 2021, 601–610)<br>(White et al. 2014, S7–S15)<br>(Xu et al. 2024, e2353331) |
| Insulin | (Sun et al. 2016a, 2334–2344)<br>(Kristensson et al. 2024, 856–863)<br>(Rose and Vona-Davis 2012, R225)<br>(Gunter et al. 2009b, 48–60)<br>(Javed et al. 2024, 1724–1736)<br>(Goodwin, Pamela J. et al. 2002, 42–51)<br>(Tseng 2015, 846)<br>(Kundaktepe et al. 2021a)<br>(Goodwin, P. J. 2011, O7) |
|  | (Goodwin, Pamela J. et al. 2008, 501–505) |
| HOMA | (Kundaktepe et al. 2021b)<br>(Pan et al. 2020, 3638–3647)<br>(Sun et al. 2016b, 2334–2344)<br>(Srinivasan et al. 2022, e21712)<br>(Igwe et al. 2015)<br>(Goodwin, P. J. 2011, O7) |
| Resistin | (Patricio et al. 2018, 29)<br>(Lee, Jung Ok et al. 2016, 18923)<br>(Wang et al. 2021, 229–239)<br>(Wang et al. 2022, 15437)<br>(Deb et al. 2021, 101178) |

**Supplemental Table 4.** Summary of colliders, mediators, and confounders identified in the directed acyclic graph (DAG) produced by DeepCEF for the breast cancer dataset.

| Confounders | Mediators | Colliders |
| --- | --- | --- |
| Breast cancer<-Resistin->HOMA | MCP.1->Resistin-> Breast cancer | MCP.1->Resistin<-Adiponectin |
| Resistin<-MCP.1->HOMA | MCP.1->Resistin->HOMA | MCP.1->Resistin<-Leptin |
| Resistin<-MCP.1->Insulin | Adiponectin->Resistin-> Breast cancer | MCP.1->Resistin<-Insulin |
| HOMA<-MCP.1->Insulin | Adiponectin->Resistin->HOMA | MCP.1->Resistin<-BMI |
| Resistin<-Adiponectin->Age | Leptin->Resistin-> Breast cancer | Adiponectin->Resistin<-Leptin |
| MCP.1<-Leptin->Resistin | Leptin->Resistin->HOMA | Adiponectin->Resistin<-Insulin |
| MCP.1<-Leptin->HOMA | Insulin->Resistin-> Breast cancer | Adiponectin->Resistin<-BMI |
| MCP.1<-Leptin->Insulin | Insulin->Resistin->HOMA | Leptin->Resistin<-Insulin |
| Resistin<-Leptin->HOMA | BMI->Resistin-> Breast cancer | Leptin->Resistin<-BMI |
| Resistin<-Leptin->Insulin | BMI->Resistin->HOMA | Insulin->Resistin<-BMI |
| HOMA<-Leptin->Insulin | Leptin->MCP.1->Resistin | Resistin-> Breast cancer<-HOMA |
| Breast cancer<-HOMA->Glucose | Leptin->MCP.1->HOMA | Resistin-> Breast cancer<-Insulin |
| Breast cancer<-Insulin->Resistin | Leptin->MCP.1->Insulin | Resistin-> Breast cancer<-Glucose |
| Breast cancer<-Insulin->HOMA | BMI->MCP.1->Resistin | Resistin-> Breast cancer<-BMI |
| Breast cancer<-Insulin->Glucose | BMI->MCP.1->HOMA | Resistin-> Breast cancer<-Age |
| Resistin<-Insulin->HOMA | BMI->MCP.1->Insulin | HOMA-> Breast cancer<-Insulin |
| Resistin<-Insulin->Glucose | BMI->Adiponectin->Resistin | HOMA-> Breast cancer<-Glucose |
| HOMA<-Insulin->Glucose | BMI->Adiponectin->Age | HOMA-> Breast cancer<-BMI |
| Breast cancer<-BMI->MCP.1 | BMI->Leptin->MCP.1 | HOMA-> Breast cancer<-Age |
| Breast cancer<-BMI->Resistin | BMI->Leptin->Resistin | Insulin-> Breast cancer<-Glucose |
| Breast cancer<-BMI->Adiponectin | BMI->Leptin->HOMA | Insulin-> Breast cancer<-BMI |
| Breast cancer<-BMI->Leptin | BMI->Leptin->Insulin | Insulin-> Breast cancer<-Age |
| Breast cancer<-BMI->HOMA | Resistin->HOMA-> Breast cancer | Glucose-> Breast cancer<-BMI |
| Breast cancer<-BMI->Insulin | Resistin->HOMA->Glucose | Glucose-> Breast cancer<-Age |
| Breast cancer<-BMI->Glucose | MCP.1->HOMA-> Breast cancer | BMI-> Breast cancer<-Age |
| Breast cancer<-BMI->Age | MCP.1->HOMA->Glucose | Leptin->MCP.1<-BMI |
| MCP.1<-BMI->Resistin | Leptin->HOMA-> Breast cancer | Resistin->HOMA<-MCP.1 |
| MCP.1<-BMI->Adiponectin | Leptin->HOMA->Glucose | Resistin->HOMA<-Leptin |
| MCP.1<-BMI->Leptin | Insulin->HOMA-> Breast cancer | Resistin->HOMA<-Insulin |
| MCP.1<-BMI->HOMA | Insulin->HOMA->Glucose | Resistin->HOMA<-BMI |
| MCP.1<-BMI->Insulin | BMI->HOMA-> Breast cancer | MCP.1->HOMA<-Leptin |
| MCP.1<-BMI->Glucose | BMI->HOMA->Glucose | MCP.1->HOMA<-Insulin |
| MCP.1<-BMI->Age | MCP.1->Insulin-> Breast cancer | MCP.1->HOMA<-BMI |
| Resistin<-BMI->Adiponectin | MCP.1->Insulin->Resistin | Leptin->HOMA<-Insulin |
| Resistin<-BMI->Leptin | MCP.1->Insulin->HOMA | Leptin->HOMA<-BMI |
| Resistin<-BMI->HOMA | MCP.1->Insulin->Glucose | Insulin->HOMA<-BMI |
| Resistin<-BMI->Insulin | Leptin->Insulin-> Breast cancer | MCP.1->Insulin<-Leptin |
| Resistin<-BMI->Glucose | Leptin->Insulin->Resistin | MCP.1->Insulin<-BMI |
| Resistin<-BMI->Age | Leptin->Insulin->HOMA | Leptin->Insulin<-BMI |
| Adiponectin<-BMI->Leptin | Leptin->Insulin->Glucose | HOMA->Glucose<-Insulin |
| Adiponectin<-BMI->HOMA | BMI->Insulin-> Breast cancer | HOMA->Glucose<-BMI |
| Adiponectin<-BMI->Insulin | BMI->Insulin->Resistin | Insulin->Glucose<-BMI |
| Adiponectin<-BMI->Glucose | BMI->Insulin->HOMA | Adiponectin->Age<-BMI |
| Adiponectin<-BMI->Age | BMI->Insulin->Glucose |  |
| Leptin<-BMI->HOMA | HOMA->Glucose-> Breast cancer |  |
| Leptin<-BMI->Insulin | Insulin->Glucose-> Breast cancer |  |
| Leptin<-BMI->Glucose | BMI->Glucose-> Breast cancer |  |
| Leptin<-BMI->Age | Adiponectin->Age-> Breast cancer |  |
| HOMA<-BMI->Insulin | BMI->Age-> Breast cancer |  |
| HOMA<-BMI->Glucose |  |  |
| HOMA<-BMI->Age |  |  |
| Insulin<-BMI->Glucose |  |  |
| Insulin<-BMI->Age |  |  |
| Glucose<-BMI->Age |  |  |

## Notes

### Competing Interest Statement

The authors have declared no competing interest.

